# Macrophage-targeted DNA methyltransferase inhibitor SGI-1027 decreases atherosclerosis in ApoE-null mice

**DOI:** 10.1101/2023.08.22.554347

**Authors:** Ana Cristina Márquez-Sánchez, Alejandro Manzanares-Guzmán, Ramón Carriles-Jaimes, Lino Sánchez-Segura, Dannia Colín-Castelán, Dan Kamen, Ekaterina K. Koltsova, Agustino Martínez-Antonio, Dalia Rodríguez-Ríos, Gloria del Carmen Romo-Morales, Gertrud Lund, Silvio Zaina

## Abstract

**Background and aims:** Correction of vascular DNA hypermethylation may slow atherogenesis. We tested the anti-inflammatory and anti-atherogenic activity of macrophage-targeted DNA methyltransferase (DNMT) inhibitor SGI-1027.

**Methods, Results:** SGI-1027 was encapsulated into human serum albumin (HSA) nanoparticles (HSANP) functionalized with the PP1 peptide, a macrophage scavenger receptor 1 ligand, fused to a FLAG epitope (S-HSANP-FLAGPP1). Nanoparticle physico-chemical characteristics predicted good marginalization towards the vascular wall, although SGI-1027 encapsulation efficiency was relatively low (∼23%). S-HSANP-FLAGPP1 were rapidly internalized compared to non-functionalized and, surprisingly, functionalized void controls, and induced a shift towards an anti-inflammatory profile of secreted cytokines in human THP-1 macrophages. S-HSANP-FLAGPP1 colonized the atheroma and induced a significant ∼44% reduction of atherosclerosis burden in the aortic tree of ApoE-null mice compared to controls. A reduction in aortic root atherosclerosis was observed, although primarily induced by HSANP irrespective of loading or functionalization. No alteration of body weight, non-vascular tissue gross histology, plasma glucose, triglyceride or cholesterol were observed. HSA whether free or structured in nanoparticles, induced a 3-4-fold increase in HDL compared to vehicle.

**Conclusions:** We confirm that DNMT inhibition is anti-atherogenic and provide proof of principle that targeted HSANP are effective carriers for those molecules. SGI-1027 displayed a novel anti-inflammatory activity that is independent of cell proliferation and therefore likely unrelated to DNMT inhibition. HDL elevation may represent an additional advantage of HSA-based nanocarriers.

## Introduction

Atherosclerosis is an inflammatory disorder of the arterial wall and an underlying cause of cardiovascular disease (CVD). Clinical complications of CVD - myocardial infarction, stroke, peripheral arterial disease - are significant causes of death and disability worldwide [1,2]. Pharmacological and lifestyle-induced reduction of hyperlipidaemia, a major risk factor for atherosclerosis, have successfully lowered the incidence of clinical complications of CVD [3]. Yet, a sizeable residual CVD risk persists after correction of hyperlipidaemia, thus underscoring the need to test candidate additional therapeutic strategies [4,5]. Modulation of gene expression by targeting the DNA methylome is among such therapeutic strategies. DNA methylation is a reversible chemical modification of the genome that participates in transcriptional regulation [6,7]. The most abundant methylated base is cytosine (5-methylcytosine) in a CpG dinucleotide context in mammalian DNA. 5-methylcytosine is produced by three catalytically active DNA methyltransferases (DNMT: DNMT1, DNMT3a and DNMT3b), whereas DNA demethylation is operated by a combination of ten-eleven translocation dioxygenases (TET1, TET2 and TET3)-catalysed 5-methylcytosine oxidation and DNA damage repair in mammals [8,9]. Complex deviations from physiological DNA methylation profiles have been observed in the atherosclerotic artery of animal models and humans. On the one hand, the DNA of affected artery is hypomethylated when compared with different vascular beds within the same individual or when the same vascular bed is compared between individuals [10–12]. On the other hand, when donor-matched and vascular bed-matched atherosclerotic and histologically normal arteries are compared, DNA hypermethylation is observed of the diseased samples [13–15]. These inconsistencies likely reflect poorly understood dynamic changes in DNA methylation during early atherogenesis [16]. Nevertheless, growing evidence indicates that DNA hypermethylation is a landmark and potential therapeutic target in atherosclerosis. Atherogenic lipoproteins and proinflammatory fatty acids induce DNA hypermethylation in cellular and animal models [17–19]. A comparison of twins discordant for CVD identified DNA hypermethylation of several relevant loci in the affected sibling [20]. Exposure to disturbed blood flow, a landmark of proatherogenic vascular microenvironment, induces DNMT expression in endothelial cells [21]. Furthermore, consistent changes were observed in the DNA demethylase TET2 in atherosclerosis: TET2 expression is decreased during neointimal formation *in vivo* and by oxidised LDL in cellular models [22,23]. Importantly, a wealth of experimental data assigns a causal role to DNA hypermethylation in atherogenesis. DNMT1 overexpression accelerates atherosclerosis in ApoE-null mice [24]. Conversely, DNMT inhibition by systemic administration of the nucleotide analogue azacytidine decreases atherosclerosis in mouse models [25–27]. Azacytidine was singled out among >3,000 drugs that promote a contractile, non-atherogenic smooth muscle cell (SMC) phenotype [28]. TET2 overexpression, which is expected to decrease global DNA methylation, is anti-atherogenic [29]. These findings underline the potential of DNMT inhibitors as therapeutics for atherosclerosis. A minimal requirement for the use of DNMT inhibitors *in vivo*, is effective targeting to the atheroma to avoid widespread DNA hypomethylation and dysregulated gene expression in non-target organs.

Here, we designed a nanoparticle (NP)-based strategy to target a DNMT inhibitor to the atheroma. We chose human serum albumin (HSA)-based NP (HSANP). HSA is an endogenous protein and building block for the synthesis of highly versatile NP with low immunogenicity that are capable of targeting macrophages [30,31]. A further reason to use HSA is to simplify the transition to future human studies. Among the available DNMT inhibitors, we chose the non-nucleoside analogue SGI-1027 (N-(4-(2-Amino-6-methylpyrimidin-4-ylamino)phenyl)-4-(quinolin-4-ylamino) benzamide; CAS no. 1020149-73-8), based on the following rationale: firstly, we avoided nucleoside analogues such as the widely used azacytidine and decitabine, as these act subsequent to incorporation into newly synthesized DNA in replicating cells, thus potentially affecting genome integrity, cell survival and plaque stability [32,33]; additionally, SGI-1027 is a suitable load for HSANP due to its hydrophobicity [30]. NP were functionalized with the short synthetic peptide PP1, a ligand of human and mouse scavenger A receptor (MSR1) for macrophage targeting [34]. We discuss our findings in the context of existing literature and the foreseeable epigenetic therapies for CVD.

## Materials and methods

### NP synthesis

HSANP were synthesized by the desolvation method as previously described [35]. Briefly, 100 mg/ml HSA (Sigma-Aldrich, no. A3782) in 10 mM NaCl, pH 9.4 was stirred using an ULTRA-TURRAX^®^ (IKA) system at 200 rpm for 30 min at room temperature. Cross-linking was carried out by adding 3 ml 4% aqueous glutaraldehyde at 0.5 ml/min rate under stirring at 150 rpm for 30 min. The resulting NP were purified by 3 cycles of differential centrifugation (10,000 rpm, 7 min). The pellet was dissolved to the original volume in 10 mM NaCl, pH 9.4. SGI-1027-loaded HSA NP (S-HSANP) were obtained by adding 1 mg/ml SGI-1027 in ethanol (1 ml/min rate) with constant stirring at 160 rpm before the cross-linking step. To determine microencapsulation and SGI-1027 loading efficiency, NP were centrifugated at 10,000 rpm for 5 min, the supernatant was discarded and replaced by the same volume of HPLC-grade absolute ethanol. Loaded SGI-1027 was extracted from NP by sonication for 15 min, pause for 30 min, further sonication for 10 min and centrifugation at 10,000 rpm for 5 min. The pellet was used to determine HSA microencapsulation efficiency and the supernatant was used to determine SGI-1027 loading efficiency and loading capacity. To measure microencapsulation efficiency, the pellet was resuspended to the original volume in ddH_2_O, transferred to a microtube of known weight, centrifugated at 10,000 rpm for 5 min and the supernatant discarded. The pellet was dried with a MAXI-dry vacuum centrifuge (Heto-Holten) at 1,300 rpm, 30°C for 45 min and weighed to obtain the total amount of HSA incorporated into NP. Microencapsulation yield was calculated as percent HSA in NP of initial amount of HSA used to prepare NP. Free SGI-1027 in the supernatant was determined by HPLC fractionation with a ZORBAX Eclipse Plus C18 column (pore size 130 Å, 1.8 μm, 2.1 mm x 50 mm; Agilent) with a reverse phase of CH_3_OH, 0.1% trifluoroacetic acid (TFA) / H_2_O, 0.1%TFA (v/v) and UV detection at 360 nm. Loaded SGI-1027 was quantified against pure standards. Loading efficiency was calculated as percent loaded (*i.e.* total – free) of total SGI-1027. Loading capacity was the percent loaded SGI-1027 of NP weight.

### NP surface functionalization with FLAGPP1

NP were functionalized with the FLAGPP1 fusion peptide, containing in N-terminus to C-terminus order: a 3-glycine spacer, a FLAG epitope (italics) and the PP1 peptide (underlined): GGG*DYKDDDD*<u>KLSLERFLRCWSDAPA</u> (>95% purity; Biomatik) [34]. Functionalization was carried out as previously described with minimal modifications (Ulbrich et al, 2011). Briefly, 800 µl FLAGPP1 (5 mg/ml in ddH_2_O) were incubated with 200 µl N-(3-dimethylaminopropyl)-N-ethylcarbodiimide in the dark for 15 min at 24°C under constant shaking at 700 rpm on a benchtop mixer. One ml NP was added, and shaking was continued for 1 h. The reaction was stopped by centrifugation at 300 rpm for 5 min. NP were purified by 3 cycles of centrifugation (10,000 rpm, 7 min), resuspended to the original volume in ddH_2_O. Functionalization was verified by immunogold labelling with InnovaCoat GOLD (Expedeon, 40nm). The conjugation solution was prepared according to the manufacturer’s instructions with the anti-FLAG Monoclonal Antibody (FG4R) at 1:3,000 dilution (ThermoFisher Scientific, no. MA1-91878). One ml functionalized NP was centrifugated at 10,000 rpm for 5 min. The pellet was resuspended in 1 ml conjugation solution, incubated for 30 min at 4°C under constant shaking and washed three times in distilled water by centrifugation at 10,500 rpm for 5 min. Antibody binding was revealed by transmission electron microscopy (TEM; Morgagni M-268, Philips/FEI). Samples were spotted on 200 mesh formvar/carbon coated copper grids (Ted Pella) and incubated for 10 min. Grids were dried at room temperature for 5 min. No positive contrast was used. Operating conditions were: 80 kV high voltage (EHT) in high magnification (1,000-140,000x), working pressure 5×10^-3^ Pa (5×10^-5^ Torr). Micrographs were captured in TIFF format, 1376×1032 pixels size, grayscale. In this format, 0 was assigned to black and 255 to white. Image analysis was performed in ImageJ with shape descriptors plugin. FLAGPP1-functionalized NP loaded with SGI-1027 or void are henceforth referred to as S-HSANP-FLAGPP1 or HSANP-FLAGPP1, respectively. In all cases, NP were kept in refrigeration until used.

### Physicochemical NP characterization

Morphometric analysis was performed by TEM. Seven μl resuspended NP were contrasted by adding 2.5 % uranyl acetate (Electron Microscopy Science) and incubated for 15 min. Sixty images of each NP version were analysed. Diameter, area, and perimeter were recorded. ζ-potential, polydispersity index (PI) and hydrodynamic ratio were obtained for technical triplicate and experimental duplicate, by dynamic light scattering (DLS) using a Zetasizer Nano ZS (Malvern Panalytical), at room temperature and a scattering angle of 13° after a 1:10 dilution in distilled water, pH 7.4.

### NP uptake by cultured macrophages

Typically, 1×10^6^ human THP-1 monocytes (ATCC no. TIB-202™) were cultured in 60 x15 mm cell culture dishes in RPMI medium supplemented with 10% foetal bovine serum, 1% penicillin/streptomycin, 2 mM glutamine at 37°C, 5% CO_2_. THP-1 monocytes were differentiated to macrophages by supplementing the medium with 50 ng/mL phorbol 12-myristate 13-acetate (PMA) for 4 h, followed by fresh 25 ng/mL PMA until complete differentiation (∼4 days). Cells were maintained in serum-free RPMI medium for 30 min before the uptake assay. NP were labelled with eosin Y, a high-affinity HSA binding dye HSA [36]. NP were incubated with 1 µl/ml eosin Y at 4°C for 15 min with gentle shaking, followed by three washes and centrifugation at 10,000 rpm for 5 min with distilled water. NP were sonicated for 5 min immediately before the assay. Following incubation with labelled NP, cells were washed three times with PBS, fixed with 4% paraformaldehyde for 20 minutes, and washed three times with PBS. Cell viability was determined by trypan blue exclusion. Cellular uptake of eosin Y-labelled NP was determined by multiphoton microscopy and flow cytometry. Prior to microscopy, cell membranes were stained with 5% SynaptoRed C2 (Sigma-Aldrich no. S6689) in ddH_2_O for 30 min. Several washes at room temperature were performed until no colour in solution was observed, followed by three washes in ice-cold PBS. Nuclei were stained with 1% DAPI (Sigma-Aldrich no. D9542) for 10 min. All sample staining and washes were performed under gentle agitation. Multiphoton microscopy was performed immediately following staining. Detection of eosin Y-labelled NP emission spectrum was performed using the LSM 880-NLO multiphoton microscopy system (Zeiss) equipped with Ti:Sapphire laser (Chameleon Vision II, Coherent) capable of tuning in the 690-1060 nm range. Images were acquired by separating the emission into three channels, blue or UV (371–440 nm), green/yellow (450–550 nm), and red (560–730 nm). The “lambda mode” scanning for detection of eosin Y-labelled NP spectral emission was performed by excitation at 810 nm. For SynaptoRed C2, an Argon laser was operated at 0.18% of power and 53 µm open pinhole with excitation at 488 nm and emission detection at 683 nm. Operating conditions for DAPI detection were Chameleon laser operated at 1.0% power with excitation at 710 nm and emission detection at 355 nm. Observation was carried out with immersion objective 60X/1.3, NA ∞-0.17, Plan-Neofluar (Zeiss).

For flow cytometry, NP-stimulated cells were harvested with 1 ml PBS, 5 mM EDTA at 4°C for 10 min with shaking. Cells were transferred to an Eppendorf tube and fixed with 2% paraformaldehyde for 15 min at room temperature. After three washes with PBS by centrifugation at 3,500 rpm for 5 min, cells were resuspended in PBS and kept in the dark until analysed. Labelled cells were detected using a 450-580nm excitation-emission wave in a FACSCanto™ II (BD Bioscience) flow cytometer. The FlowJo software (v10.4.1) was used for analysis.

### Cytokine determination in cell culture medium

IL-1ß, IL-6, IL-8, IL-10 and INFα polypeptides were determined by iqPCR (PreciseDX Lab, Chihuahua, Chi., Mexico). Briefly, 200 µl medium were used for the analyte capture, using a 1:10 dilution of monoclonal antibodies (Santa Cruz Biotechnology): anti-IL-1ß (H-153; sc-7884), anti-IL-6 (E-4; sc-28343), anti-IL-8 (C-11; sc-376750), anti-IL-10 (E-10; sc-8438), anti-INFα (FL-189; sc-20106). After a 15 min incubation at room temperature, 400 µl PBS were added. Samples were centrifuged at 10,000 rpm for 5 min, the supernatant was discarded, and a secondary antibody coupled to a synthetic DNA fragment was added following manufacturer’s instructions (Thunder-Link Plus Oligo Conjugation System, Expedeon). After 2 washing cycles, qPCR was performed, using Phire Hot Start II DNA Polymerase Master Mix Green (F126L, Thermo Scientific) following a 3-step reaction cycle: initial denaturation at 98 °C, 30 s; 30 cycles: 98 °C for 5 s, 64 °C for 5 s, 72 °C for 5s; final extension at 72 °C for 60 s. Samples were quantified in triplicate.

### Mouse procedures

The laboratory animal protocol was approved by the committee for ethics in research of the University of Guanajuato (approval no. CIBIUG-P10-2017) and performed in agreement with the official Mexican guideline for the use of laboratory animals NOM-062-ZOO-1999.

In preliminary experiments aimed to detect subcutaneously injected HSANP in peripheral blood, whole tail blood and Tris-buffered saline (TBS) with 2% Tween (TBS-T) (2.5 µl each) were mixed and carefully spotted on a nitrocellulose membrane. After drying at room temperature, equal loading was estimated by Ponceau red staining. Membranes were blocked with TBS-T, 5% fat-free milk for 1.5 h. After incubation with a 1:2,500 dilution of an anti-HSA, HRP-conjugated monoclonal antibody (ab24458, Abcam) for 30 min at room temperature, the membrane was washed three times with TBS-T, 0.1% fat-free milk for 30 min at room temperature. Antibody binding was revealed with ECL^™^ Prime Western Blotting Detection (Amersham). HSA was determined by comparison with a standard curve using free HSA and the average value at time 0 was subtracted from all other time point data.

High fat diet-fed nullizygous ApoE-null mice in a C57BL/6 background, a model of hyperlipidaemia-induced atherosclerosis, were used [37]. Mice were housed in groups of 4-5 in a controlled environment with 12 h day-light cycle and *ad libitum* access to food and water. Twenty-week old female and male mice were fed for 8 weeks with a high fat diet (30% fat) that was elaborated by us as follows. A standard chow (Rodent 5001, LabDiet) was pulverized and mixed with pork lard (0.16:1 lard:chow ratio, w:w). The resulting paste was pelleted and stored at 4 °C for no longer than 5 days before use. Mice were treated with HSANP (n=20), HSANP-FLAGPP1 (n=10), S-HSANP-FLAGPP1 (n=20), vehicle (n=20), or an unstructured mix of SGI-1027 and native HSA (SGI-HSA) (n=10) by weekly injections during the 8-week high-fat diet feeding. That sample size exceeded the one used in comparable studies where a 40% reduction of the size of atheroma - deemed here as the lowest biologically acceptable effect - was observed [26]. NP were sonicated for 5 min prior to injection. Overnight-fasting mice were sacrificed 1 week after the last injection. Glucose was measured in a tail blood drop (One Touch Ultra 2, Johnson & Johnson) immediately prior to sacrifice. Sacrifice was performed by decapitation under isoflurane-induced anaesthesia. Total blood was collected at decapitation in EDTA (Vacutainer, BD) for plasma total cholesterol, triglycerides, and HDL determination by the Clinical Analysis Laboratory, Department of Medical Sciences, University of Guanajuato, following standard colorimetric methods. The aortic root, the remaining aorta (arch, thoracic, abdominal to the caudal bifurcation), liver, kidney, adipose tissue, ventricular (lower) heart portion, were dissected and processed for downstream analysis.

### Histology

Liver, kidney, adipose tissue, and ventricular heart portion were transferred to 300 µL RNAStill (RNA stabilization agent, RBC) at dissection, stored at -20°C and transferred to 1% formaldehyde. The aortic root was dissected, washed in PBS, and fixed in 1% formaldehyde. Fixation was for at least 24 h prior to paraffin embedding. Serial sections (6 µm) of the aortic root were stained with hematoxylin and eosin (HE), Masson trichrome, or periodic acid–Schiff (PAS) according to standard protocols. Three sections were used to determine cellularity, basal lamina, and collagen using a U-DO3 microscope (Olympus Optical CO, LTD); images were digitalized and examined with Image J software. For this purpose, all images were taken at 20X, and the regions of interest were traced where loss of continuity of media smooth muscle was observed. Original images were converted to 8 bits, and a threshold was obtained. This shadow was overlaid on the original image, to visually separate the cells. Cellularity and foam cell counting were limited for those with complete nuclei and identifiable cytoplasm. PAS-stained sections were used for the determination of the lesion area, using the ImageJ software version 1.52 (NIH, USA). Smooth muscle was stained with an anti-smooth muscle actin (SMA) antibody (rabbit-anti-mouse, Abcam, no. 5694). Briefly, samples were dewaxed by heating at 65 °C for 1 h, rehydrated in a xylene-ethanol train (three changes of xylene during 15 min each, one change of xylene-ethanol (1:1), two changes of 100% ethanol, one change of 95% ethanol and two changes of 70% ethanol during 5 min each), and washed with ddH2O. Antigen retrieving was performed by heating the samples in a 1X citrate solution (Electron Microscopy Sciences, no. 64142-08) at 94 °C for 45 min. After cooling, samples were washed twice with PBS for 15 min, and endogenous peroxidase was blocked by incubation in 3% H_2_O_2_. Samples were washed in PBS and then in PBS, 0.05% Triton X-100 (PBST) for 10 min. Samples were blocked with a 5% BSA solution at 4 °C for 45 min. After blocking, samples were incubated with the anti-SMA primary antibody (1:200) diluted in PBST, 1% BSA, at 4 °C overnight. Samples were rinsed twice with PBS for 15 min and with PBST for 5 min, followed by incubation with an anti-Rabbit-HRP secondary antibody (GeneTex-GTX 83399 OneStep Polymer HRP) according to manufacturer’s instructions. Staining was performed with a DAB substrate system (Abcam, no. ab64238) according to manufacturer’s instructions, followed by washing in ddH2O for 5 min. Counterstaining was performed using hematoxylin (Sigma-Aldrich, no. HHS32) and a final incubation in 0.25% ammonium water, followed by dehydration (70% ethanol for 5 min, three times; 100% ethanol for 5 min, two times; and three changes of xylene for 5 min). Samples were mounted using Permount mounting medium solution (Fisher Chemical, SP15). All images were acquired with a Nikon Eclipse 80i microscope or EVOS Auto FL2. Liver, kidney, and heart tissue sections (7 µm) were stained with HE and examined by a trained pathologist. For the assessment of the aortic lesion area, whole mount oil red O (ORO) staining was performed essentially as described [38]. Images were digitalized and the lesion area was calculated using the ImageJ software. The lesion area was expressed as a percentage of the area of lesions (dark red), relative to the total surface area of the aortic lumen. For NP detection in the aorta, 6 µm paraffin sections of the aortic root of mice treated with vehicle or S-HSANP-FLAGPP1 were deparaffinized and prepared as stated for IHC. We used immunogold for the detection of the FLAG epitope in S-HSANP-FLAGPP1, using an anti-FLAG antibody coupled to 40 nm gold nanoparticles. Immunogold was prepared as described in the previous section and incubated with the tissue overnight at 4 °C. After three 10-minute washes with PBS, samples were stained with DAPI (1:1000, 20 min), followed by two washes with PBS. Samples were mounted using Permount mounting media. Images were acquired in a Leica Stellaris 8-STED Confocal Microscope (60 X oil-objective) by separating the emission in two channels, blue for DAPI (400–450 nm), and red for gold (590–650 nm). Gold-labelled NP were excited at 530 nm. At the sites with signal detection, zoom-in (3.0) was performed, and 14 consecutive images were acquired along the Z-axis. A 3D image of the signal detected was extracted from one of these Z-stacks. Images were collected and further analyzed with the FIJI software. Contrast and colour adjustment were performed using the same parameters across all images.

### Statistics

The Kruskal Wallis test with Bonferroni *post hoc* or Mann-Whitney *U* test were used for non-normally distributed data. Paired Student’s *t* test and ANOVA were used in the case of normal distribution. Significance threshold was set at *p=*0.05 (two-tailed). Tests were performed with SPSS v20 (IBM).

## Results

### NP physico-chemical properties

TEM and DLS analysis showed the following characteristics for HSANP: ∼300 nm or ∼100 nm diameter, as determined by TEM or DLS analysis, respectively; acceptable size homogeneity; markedly negative surface charge (**Table 1**). Loading with SGI-1027 did not significantly affect any of those parameters (see **Supplementary Figure 1** for representative HPLC chromatograms). Functionalization with FLAGPP1 significantly increased NP hydrodynamic diameter by ∼3-fold when HSANP and S-HSANP-FLAPP1 were compared (*p*<0.05) and promoted heterogeneity in NP size and ζ-potential (*p*<0.05). Representative TEM images of functionalized and control NP are shown in **Supplementary Figure 2**. HPLC-assessed efficiency of SGI-1027 loading onto S-HSANP-FLAGPP1 was ∼23%. As for S-HSANP-FLAGPP1 composition, the SGI-1027:HSA ratio (µg:g) was ∼0.9.

**Table 1.** NP physicochemical characteristics.

|  | HSANP | S-HSANP |
| --- | --- | --- |
| Diameter (nm)<sup>a</sup> | 308.3 ± 106.8 | 311.7 ± 109.7 |
| Roundness<sup>a</sup> | 0.8 ± 0.2 | 0.8 ± 0.1 |
| Shape factor<sup>a</sup> | 0.5 ± 0.1 | 0.4 ± 0.1 |
| Hydrodynamic diameter (nm)<sup>b</sup> |  |  |
| Not functionalized | 104.2 ± 14.4 | 141.8 ± 4.6 |
| Functionalized with FLAGPP1 | 296.4 ± 72.6 | 447.1 ± 131.2 |
| Polydispersity index<sup>b</sup> |  |  |
| Not functionalized | 0.5 ± 0.1 | 0.6 ± 0.1 |
| Functionalized with FLAGPP1 | 0.1 ± 0.1 | 0.9 ± 0.1* |
| ζ-potential<sup>b</sup> |  |  |
| Not functionalized | -43.7 ± 0.5 | -42.2 ± 0.3 |
| Functionalized with FLAGPP1 | -39.5 ± 1.3 | -44.6 ± 0.8* |
| Microencapsulation efficiency (%)<sup>c</sup> | 55.5 | 56.0 |
| Loading efficiency (%)<sup>c</sup> | not applicable | 22.6 |

Values are average ± SD. HSANP and S-HSANP, void and SGI-1027-loaded NP, respectively. <sup>a</sup>n=60 individual NP per group. <sup>b</sup>means of technical triplicate measure; <sup>c</sup>mean from a technical duplicate measure. Significance is relative to HSANP. \*, $p<0.05$ , Kruskal-Wallis test.

### Effects of NP in cultured THP-1 macrophages

No significant effect of NP on cell viability was observed (**Supplementary Figure 3**). Flow cytometry and fluorescence microscopy analysis of NP-challenged THP-1 macrophages revealed a marked, incubation time-dependent uptake of FLAGPP1-functionalized NP in the order S-HSANP-FLAGPP1 > HSANP-FLAGPP1, with readily detectable uptake of S-HSANP-FLAGPP1 at the 15 min timepoint. By contrast, HSANP uptake was comparatively slow (**Figure 1A,B; Supplementary Figure 4**). Internalized NP were mostly in cytoplasmic droplet-like structures, although optical section data were consistent with nuclear localization of a subset of S-HSANP-FLAGPP1 (**Figure 1C**).

**Figure 1.**
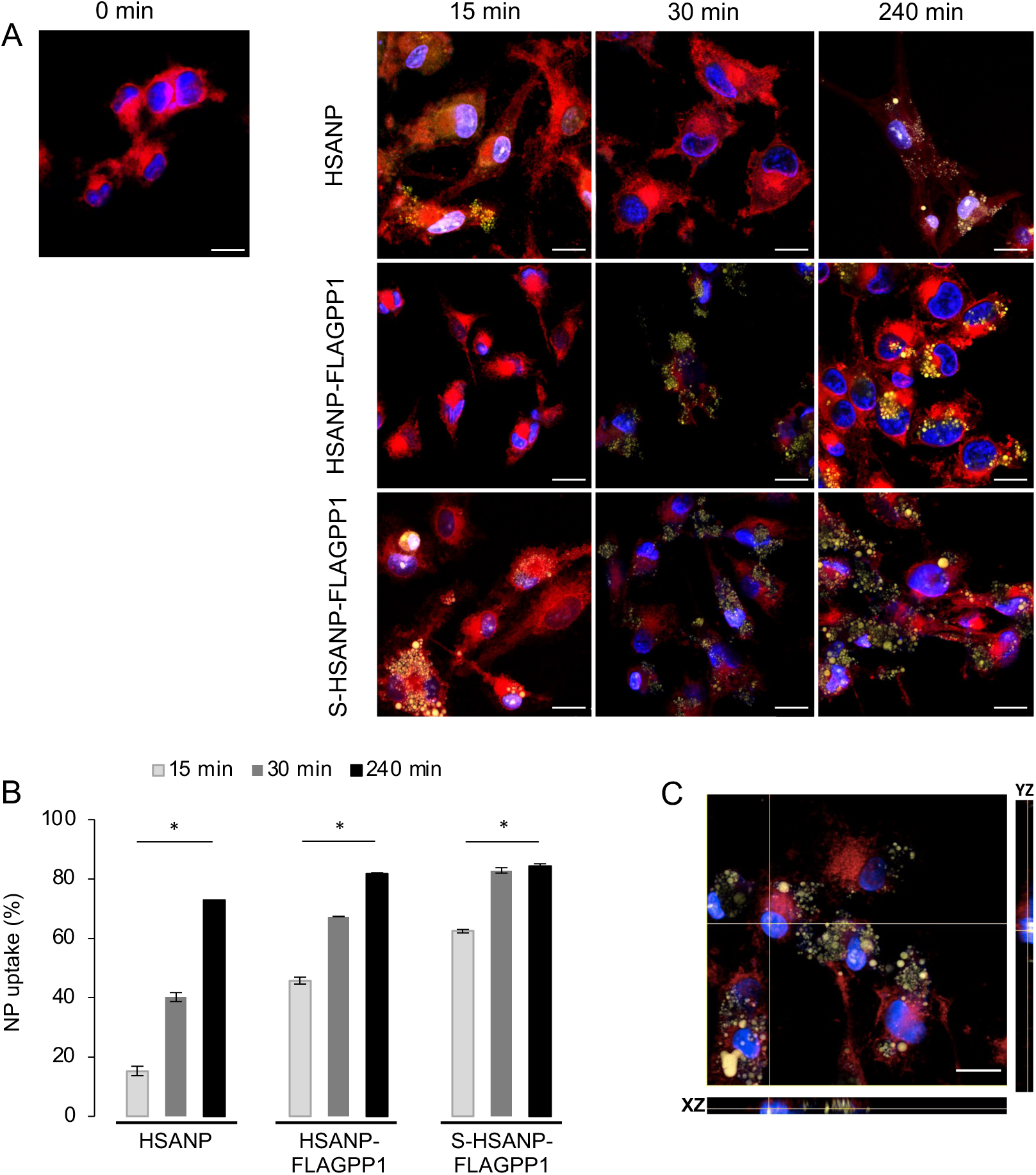
NP uptake by THP-1 macrophages. A, representative images of fluorescence microscopy analysis of cells challenged for the indicated time lengths. Blue, DAPI (nuclei); red, SynaptoRed C2 (cell membranes); yellow, eosin Y-labelled NP. Bar, 16 µm. B, flow cytometry-based analysis of NP uptake by THP-1 macrophages treated as in A. Notice the comparatively faster S-HSANP-FLAGPP1 uptake (n=3). Kruskal-Wallis test with paired-comparisons *post hoc* test; *, *p*<0.05. C, confocal microscopy-generated single-cell orthogonal view of eosin Y fluorescence. Notice the presence of eosin Y fluorescence within the nucleus. Bar, 16 µm. HSANP, void NP; HSANP-FLAGPP1, functionalized void NP; S-HSANP-FLAGPP1, SGI-1027-loaded and functionalized NP.

NP elicited distinct effects on the secreted cytokine profile. HSANP-FLAGPP1 and HSANP induced weak changes, whereas S-HSANP-FLAGPP1 modestly increased IL-10 (∼50%), but markedly (2-4-fold) augmented IL-1ß, IL-8 and INFα in a time-dependent fashion (**Figure 2**). S-HSANP-FLAGPP1 induced a 2.7-fold higher IL-10/IL-1ß ratio compared to HSANP-FLAGPP1 or HSANP, suggestive of a net anti-inflammatory effect (**Figure 2**).

**Figure 2.**
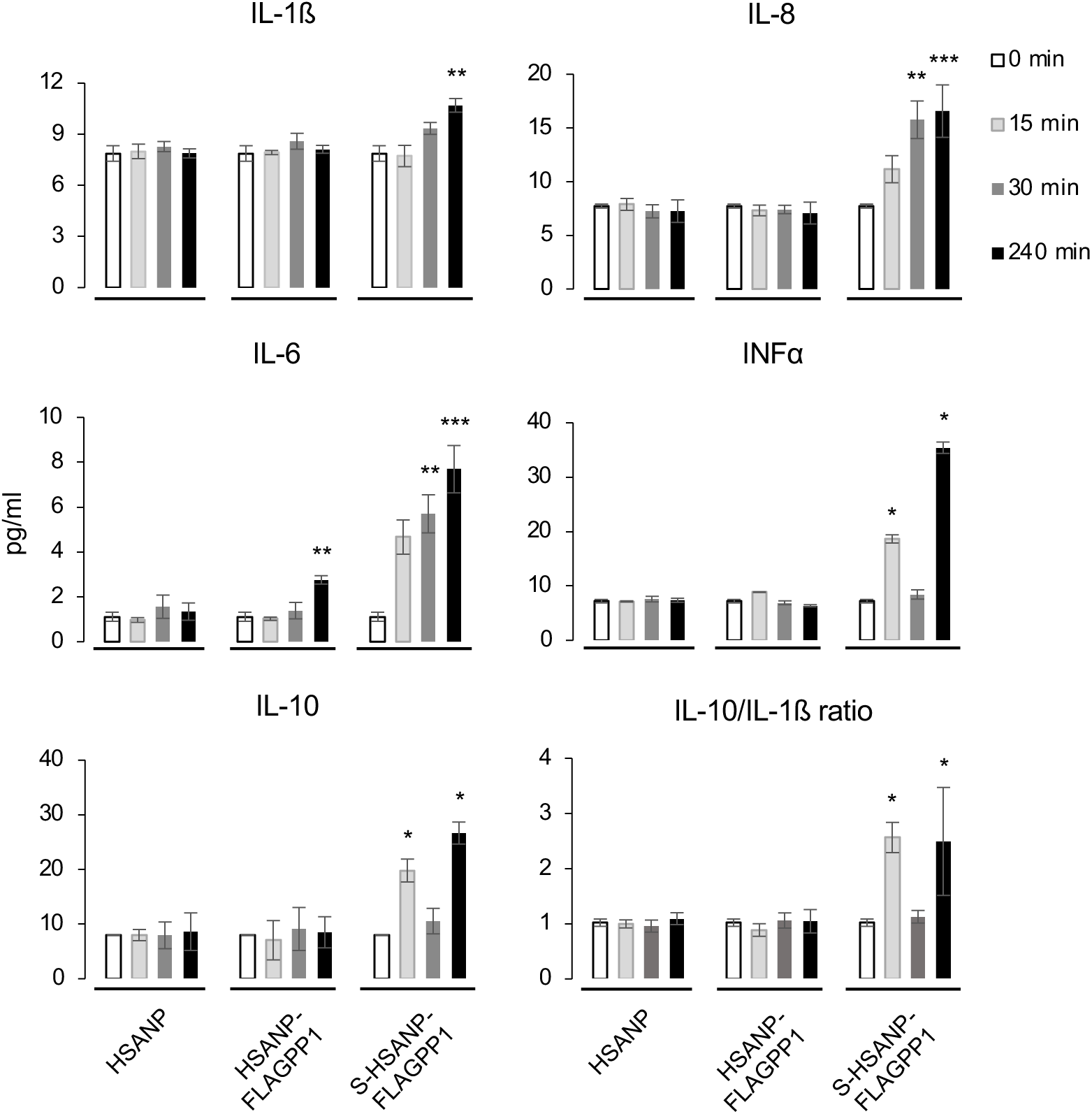
NP-induced modulation of secreted cytokine profile in THP-1 macrophages. Data are average±SD. HSANP, void NP; HSANP-FLAGPP1, functionalized void NP; S-HSANP-FLAGPP1, SGI-1027-loaded and functionalized NP. n=3. Significance of comparisons with the 0 min time point is shown. Kruskal-Wallis test, paired-comparisons-Bonferroni ajusted *post hoc* test. *, *p*<0.05, **, *p*<0.01, ***, *p*<0.001

### Feasibility of the subcutaneous route for *in vivo* administration of HSANP

Since repeated injection into the tail vein poses ethical and practical issues in dark-skinned mice, we first assessed alternative administration routes. Intraperitoneal administration was discarded based on concerns that peritoneal macrophages would sequester FLAGPP1-functionalized NP. Therefore, we tested the feasibility of the subcutaneous route. We were encouraged by the successful reduction of intimal hyperplasia by subcutaneously injected polymeric NP [39]. Six-week old female C57BL/6 mice were injected below the dorsal neck skin with 150 µl HSANP (1 mg protein/ml in PBS) or PBS (n=3 each). Fifty µl tail tip peripheral blood were obtained immediately before the injection (time 0) and at four time points up to 48 h post-injection. A significant increase in HSA with post-injection time was detected in peripheral blood (*p*=2.3×10^-4^, ANOVA and Bonferroni *post hoc* test) (**Supplementary Figure 5A**). Detected HSA at 48 h corresponded to ∼20% of the total injected NP.

### Gross *in vivo* effects of NP

We designed a weekly, single-dose protocol for *in vivo* administration of NP to ApoE-null mice. The protocol aimed to achieve a 7 µM SGI-1027 concentration in the total peripheral blood volume calculated based on recipient mouse body weights. That SGI-1027 concentration is within the reported 6-13 µM IC_50_ range and below the 20 µM toxic dose observed in rat hepatocarcinoma cells [40–42]. A schematic view of the protocol of ApoE-null mice treatment with NP is shown in **Figure 3A**.

**Figure 3.**
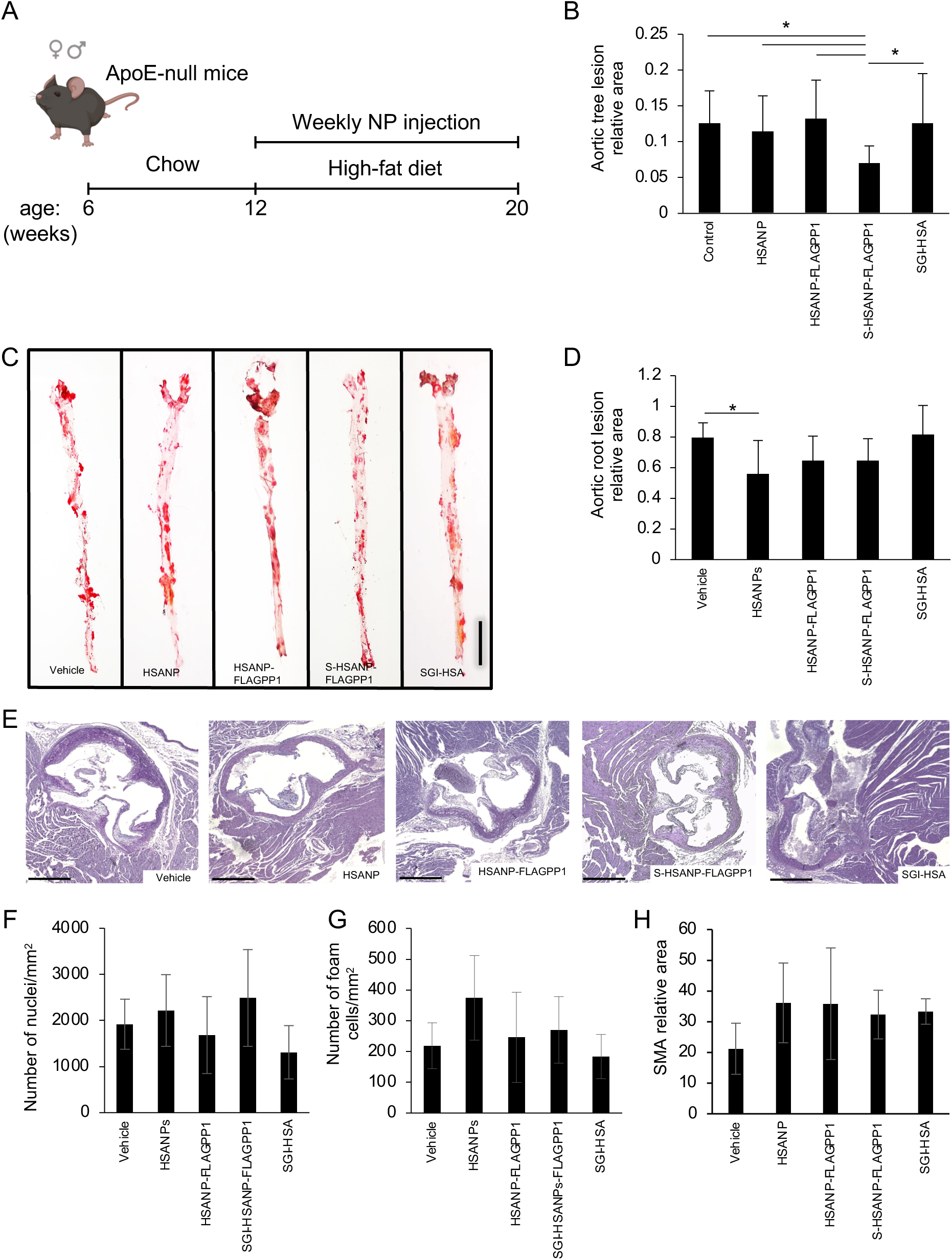
Effects of NP administration *in vivo*. A, schematic view of the experimental design of ApoE-null mice treatment with NP (mouse drawing: BioRender). B,C, quantitative analysis of atheroma size in whole mount aortas (n=20 per group) and aortic roots (n=10 per group), respectively. D, representative images of oil red O-stained atherosclerotic lesions in whole mount aortas. E, representative images of periodic acid-Schiff-stained atherosclerotic lesions in aortic roots. F-H, aortic root atheroma cellularity (n=10 per group), foam cell (n=10 per group) and smooth muscle actin (n=5 per group) abundance. Graph data are average±SD. Kruskal-Wallis and Bonferroni *post hoc* test. *, *p*<0.05. HSANP, void NP; HSANP-FLAGPP1, functionalized void NP; S-HSANP-FLAGPP1, SGI-1027-loaded and functionalized NP; SGI-HSA, unstructured mixture of SGI-1027 and native HSA. Bars in D, E are 5 mm and 100 µm respectively.

We first assessed the effects of NP (HSANP, HSANP-FLAGPP1, S-HSANP-FLAGPP1), and SGI-1027 mixed with free HSA (SGI-HSA) on general well-being, metabolic parameters, and non-vascular tissue gross morphology. No obvious alteration of mobility or fur appearance was detected. No significant change in body weight was observed at treatment endpoint, compared to first day of treatment (**Table 2**). In one case, a tangible lump was detected at the injection site. The treatment did not elicit any statistically significant change in plasma glucose, total cholesterol, or triglycerides. Conversely, all treatments induced a ∼3-4-fold increase in plasma HDL relative to vehicle, although differences reached statistical significance only for HSANP-FLAGPP1 and SGI-HSA.

**Table 2.**
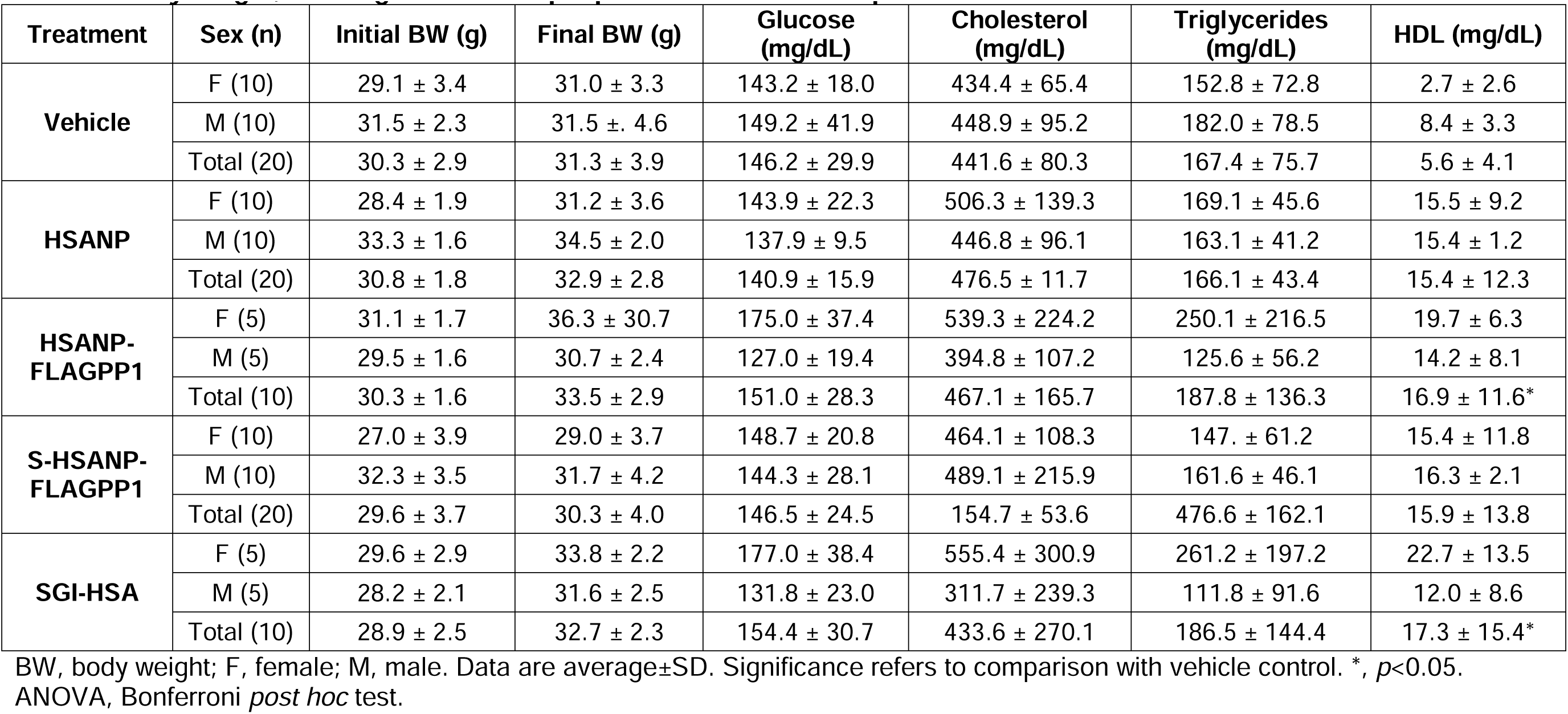
Body weight, serum glucose and lipid profile of NP-treated ApoE-null mice.

| Treatment | Sex (n) | Initial BW (g) | Final BW (g) | Glucose (mg/dL) | Cholesterol (mg/dL) | Triglycerides (mg/dL) | HDL (mg/dL) |
| --- | --- | --- | --- | --- | --- | --- | --- |
| **Vehicle** | F (10) | 29.1 ± 3.4 | 31.0 ± 3.3 | 143.2 ± 18.0 | 434.4 ± 65.4 | 152.8 ± 72.8 | 2.7 ± 2.6 |
|  | M (10) | 31.5 ± 2.3 | 31.5 ± 4.6 | 149.2 ± 41.9 | 448.9 ± 95.2 | 182.0 ± 78.5 | 8.4 ± 3.3 |
|  | Total (20) | 30.3 ± 2.9 | 31.3 ± 3.9 | 146.2 ± 29.9 | 441.6 ± 80.3 | 167.4 ± 75.7 | 5.6 ± 4.1 |
| **HSANP** | F (10) | 28.4 ± 1.9 | 31.2 ± 3.6 | 143.9 ± 22.3 | 506.3 ± 139.3 | 169.1 ± 45.6 | 15.5 ± 9.2 |
|  | M (10) | 33.3 ± 1.6 | 34.5 ± 2.0 | 137.9 ± 9.5 | 446.8 ± 96.1 | 163.1 ± 41.2 | 15.4 ± 1.2 |
|  | Total (20) | 30.8 ± 1.8 | 32.9 ± 2.8 | 140.9 ± 15.9 | 476.5 ± 11.7 | 166.1 ± 43.4 | 15.4 ± 12.3 |
| **HSANP-FLAGPP1** | F (5) | 31.1 ± 1.7 | 36.3 ± 30.7 | 175.0 ± 37.4 | 539.3 ± 224.2 | 250.1 ± 216.5 | 19.7 ± 6.3 |
|  | M (5) | 29.5 ± 1.6 | 30.7 ± 2.4 | 127.0 ± 19.4 | 394.8 ± 107.2 | 125.6 ± 56.2 | 14.2 ± 8.1 |
|  | Total (10) | 30.3 ± 1.6 | 33.5 ± 2.9 | 151.0 ± 28.3 | 467.1 ± 165.7 | 187.8 ± 136.3 | 16.9 ± 11.6* |
| **S-HSANP-FLAGPP1** | F (10) | 27.0 ± 3.9 | 29.0 ± 3.7 | 148.7 ± 20.8 | 464.1 ± 108.3 | 147. ± 61.2 | 15.4 ± 11.8 |
|  | M (10) | 32.3 ± 3.5 | 31.7 ± 4.2 | 144.3 ± 28.1 | 489.1 ± 215.9 | 161.6 ± 46.1 | 16.3 ± 2.1 |
|  | Total (20) | 29.6 ± 3.7 | 30.3 ± 4.0 | 146.5 ± 24.5 | 154.7 ± 53.6 | 476.6 ± 162.1 | 15.9 ± 13.8 |
| **SGI-HSA** | F (5) | 29.6 ± 2.9 | 33.8 ± 2.2 | 177.0 ± 38.4 | 555.4 ± 300.9 | 261.2 ± 197.2 | 22.7 ± 13.5 |
|  | M (5) | 28.2 ± 2.1 | 31.6 ± 2.5 | 131.8 ± 23.0 | 311.7 ± 239.3 | 111.8 ± 91.6 | 12.0 ± 8.6 |
|  | Total (10) | 28.9 ± 2.5 | 32.7 ± 2.3 | 154.4 ± 30.7 | 433.6 ± 270.1 | 186.5 ± 144.4 | 17.3 ± 15.4* |

BW, body weight; F, female; M, male. Data are average±SD. Significance refers to comparison with vehicle control. \*, $p < 0.05$ . ANOVA, Bonferroni *post hoc* test.

We also estimated the ability of NP to accumulate in the atheroma and off-target, *i.e.* in non-vascular bed tissue. FLAGPP1-functionalized, but not unfunctionalized, NP could be detected in the aortic atheroma, particularly at the edge of the lesion (**Supplementary Figure 5B-D**). As for off-target effects, we focused on tissue gross morphology, as assessed by HE staining. Liver, kidney or heart morphology was not detectably altered (**Supplementary Figure 6**).

### NP mitigated atherosclerosis in ApoE-null mice

Atherosclerosis was quantified in whole mount aortic arch and descending aorta, and in aortic root sections. S-HSANP-FLAGPP1 elicited a marked reduction of lesion occupancy of the aortic arch and descending aorta compared to vehicle (∼44%, *p*<0.001) and to all other treatments (**Figure 3B,C**). In the aortic root, S-HSANP-FLAGPP1 reduced lesion size by ∼40% (*p*<0.05), in comparison with control mice (**Figure 3D,E**). HSANP and HSANP-FLAGPP1 elicited similar effects (∼44% and ∼35%, respectively). Aortic root plaque cellularity, number of foam cells or SMA abundance were not significantly different from vehicle controls in any of the treatments (**Figure 3F-H**).

## Discussion

We report for the first time the synthesis of NP loaded with a non-nucleoside analogue DNMT inhibitor and functionalized with a MSR1 peptidic ligand. Structurally, the physico-chemical properties of HSANP predict good *in vivo* stability and marginalization - *i.e.* the ability to adhere to the vascular wall: non-neutral charge, µm-region size and imperfectly spherical shape [43,44]. Functionally, those NP exerted anti-inflammatory and anti-atherogenic effects in cultured human THP-1 macrophages and *in vivo* in ApoE-null mice. Our findings provide several novel insights. First, we confirm previous reports that DNMT inhibitors are anti-atherogenic [26,45]. Yet, although DNA demethylation may contribute to the effects of HSA NP *in vivo*, the relatively rapid anti-inflammatory response observed in non-replicating macrophages hints at a novel activity of SGI-1027, independent of inhibition of *de novo* DNA methylation. SGI-1027 increased secretion of the potent anti-inflammatory cytokine IL-10, the pleiotropic IL-6, and the ratio between IL-10 and IL-1ß - an inflammatory cytokine. IL-10 decreases the proinflammatory response in vascular lesions, preventing intimal hyperplasia by inhibiting the activity of the NFl1lB transcription factor, the chemokine MCP1, and the growth factor bFGF [46–48]; IL-10 also reduces the expression of matrix metalloproteinases, modulates lipid metabolism in macrophages and limits monocyte recruitment to the plaque [49–51]. Taken together, our data suggest that SGI-1027 may represent a previously unappreciated double-edged tool that targets both the DNA methylome and the macrophage inflammatory status, perhaps by rapid transcriptional regulation or biochemical modulation of cytokine translation and secretion. Further studies are warranted to clarify the molecular mechanisms underlying that response. Notably, the nucleoside analogue DNMT inhibitor azacytidine can elicit responses that are independent of DNMT inhibition [52]. Also, we confirm the ability of the PP1 peptide to be internalized by human macrophages and to colonize the murine atheroma [34]. Interestingly, our data indicate that loaded SGI-1027 increases PP1-mediated uptake, possibly by conferring physico-chemical properties to NP that facilitate PP1 binding to MSR1 or promote receptor-independent uptake.

The extent of decrease of the aortic tree lesion burden (∼44%) is comparable to results of other studies using DNMT inhibitors, such as azacytidine [26,45] although the treatment period was shorter in our study (8 weeks compared to 30 and 12 weeks, respectively). Other studies using chemotherapeutic agents such as paclitaxel or methotrexate showed a comparable decrease of lesion area [53]. The *in vivo* anti-atherogenic effects of FLAGPP1-functionalized, SGI-1027-loaded NP were robust in the descending aorta, but less selective for NP composition in the aortic root. The data point to heterogeneity of ApoE-null atheromas across different locations along the aorta. To our knowledge, evidence for such differences is anecdotal and has not been systematically investigated. Interestingly, the observation that HSANP irrespective of payload or functionalization, but not unstructured HSA, tend to decrease the size of aortic root atheromas, suggests an unexpected anti-atherogenic activity of HSANP *per se*. Based on the ability of serum albumin to avidly bind fatty acids, the anti-atherogenic activity of HSANP may be due to increased lipid sponging by HSA aggregates compared to individual HSA polypeptides, resulting in favourable changes in cellular free lipid pool within the atheroma. Our data show that the anti-atherogenic activity of HSANP is the result of proportionate deceleration of the expansion of the atheroma, rather than targeted reduction of any of its individual components - cellularity, foam cells, smooth muscle. That outcome is reassuring from the viewpoint of prospective translational applications, as it suggests that a reduction of the atheroma burden is achieved without compromising the stability of the lesion. An additional layer of the anti-atherogenic effect of HSANP is plasma HDL reduction. That effect is elicited by HSA *per se*, irrespective of whether structured in NP or free. The data echo evidence that hypoalbuminaemia is a proposed predictor of CVD [54] and serum albumin is directly associated with HDL [55,56]. Our model of serum albumin perfusion may help to understand the functional relationship of hypoalbuminaemia with lipoprotein metabolism and CVD risk.

As for off-target effects, the absence of any change in body weight, homeostasis of plasma glucose, cholesterol or triglycerides, or any gross histological morphology of non-vascular bed tissues, suggests that HSANP can deliver their cargo specifically to target organs without inducing any systemic gross metabolic alterations.

Among limitations of our work, is the use of glutaraldehyde as cross-linker, which limits future translational applications. Furthermore, the efficiency of SGI-1027 loading needs improving. Also, we could not obtain any detailed information on the effects of NP on the vascular DNA methylome and gene expression. Critically, longer duration of treatment is necessary to fully appreciate tolerance, toxicity and any undesired secondary effect.

In conclusion, despite the aforementioned limitations, our study provides encouraging proof of concept that treatment with NP-loaded DNMT inhibitors may be a valuable strategy to prevent or treat atherosclerosis. Also, we demonstrate a novel, likely independent of DNMT inhibition, anti-inflammatory activity of SGI-1027. Future research should focus on NP with amenable physico-chemical characteristics for human studies.

## Author contributions

ACM-S carried out most of the experimental work, discussed results, suggested new experiments, and wrote the first manuscript draft. AM-S performed early work on NP synthesis and administration route in mice. RC-J advised and supervised NP characterization. LS-S advised and supervised NP synthesis. DC-C, DK, AM-A, DR-R and GdCR-M supported experimental work. EKK critically revised the data. GL participated in the study design and critically revised the data. SZ designed and supervised the study, and prepared the final manuscript version.

## Acknowledgements

We thank the personnel of Cedars-Sinai Medical Center Microscopy Core, Los Angeles, CA, USA; Dr. Jiani Zhu, Postdoctoral Scientist, Medicine Department, Cedars Sinai Medical Center, Los Angeles, CA, USA; and the personnel of the Flow Cytometry Core, XXI Century Medical National Center, Mexico City, Mexico, for their generous assistance.

## Footnotes

§ In memory of Dr. Jorge A. Martínez García, excellent colleague and pathologist, generous collaborator.

## References

[1] C.D. Mathers, D. Loncar, Projections of global mortality and burden of disease from 2002 to 2030, PLoS Med. 3 (2006) e442. 10.1371/journal.pmed.0030442.

[2] P. Song, Z. Fang, H. Wang, Y. Cai, K. Rahimi, Y. Zhu, F.G.R. Fowkes, F.J.I. Fowkes, I. Rudan, Global and regional prevalence, burden, and risk factors for carotid atherosclerosis: a systematic review, meta-analysis, and modelling study, Lancet Glob. Heal. 8 (2020) e721–e729. 10.1016/S2214-109X(20)30117-0.

[3] W. Herrington, B. Lacey, P. Sherliker, J. Armitage, S. Lewington, Epidemiology of Atherosclerosis and the Potential to Reduce the Global Burden of Atherothrombotic Disease, Circ. Res. 118 (2016) 535–546. 10.1161/CIRCRESAHA.115.307611.

[4] Y. Matsuura, J.E. Kanter, K.E. Bornfeldt, Highlighting Residual Atherosclerotic Cardiovascular Disease Risk, Arterioscler. Thromb. Vasc. Biol. 39 (2019) e1–e9. 10.1161/ATVBAHA.118.311999.

[5] N. Arnold, K. Lechner, C. Waldeyer, M.D. Shapiro, W. Koenig, Inflammation and Cardiovascular Disease: The Future, Eur. Cardiol. Rev. 16 (2021) e20. 10.15420/ecr.2020.50.

[6] M.B. Stadler, R. Murr, L. Burger, R. Ivanek, F. Lienert, A. Schöler, E. van Nimwegen, C. Wirbelauer, E.J. Oakeley, D. Gaidatzis, V.K. Tiwari, D. Schübeler, DNA-binding factors shape the mouse methylome at distal regulatory regions., Nature. 480 (2011) 490–495. 10.1038/nature10716.

[7] T.M. Bertozzi, N. Takahashi, G. Hanin, A. Kazachenka, A.C. Ferguson-Smith, A spontaneous genetically-induced epiallele at a retrotransposon shapes host genome function, Elife. 10 (2021). 10.7554/eLife.65233.

[8] S. Seisenberger, J.R. Peat, T.A. Hore, F. Santos, W. Dean, W. Reik, Reprogramming DNA methylation in the mammalian life cycle: building and breaking epigenetic barriers, Philos. Trans. R. Soc. B Biol. Sci. 368 (2012) 20110330. 10.1098/rstb.2011.0330.

[9] M.V.C. Greenberg, D. Bourc’his, The diverse roles of DNA methylation in mammalian development and disease, Nat. Rev. Mol. Cell Biol. 20 (2019) 590– 607. 10.1038/s41580-019-0159-6.

[10] G. Lund, L. Andersson, M. Lauria, M. Lindholm, M.F. Fraga, A. Villar-Garea, E. Ballestar, M. Esteller, S. Zaina, DNA methylation polymorphisms precede any histological sign of atherosclerosis in mice lacking apolipoprotein E, J. Biol. Chem. 279 (2004) 29147–29154. 10.1074/jbc.M403618200.

[11] E. Aavik, H. Lumivuori, O. Leppänen, T. Wirth, S.-K. Häkkinen, J.-H. Bräsen, U. Beschorner, T. Zeller, M. Braspenning, W. van Criekinge, K. Mäkinen, S. Ylä-Herttuala, Global DNA methylation analysis of human atherosclerotic plaques reveals extensive genomic hypomethylation and reactivation at imprinted locus 14q32 involving induction of a miRNA cluster, Eur. Heart J. 36 (2014) 993–1000. http://eurheartj.oxfordjournals.org/content/early/2014/11/14/eurheartj.ehu437.abstract.

[12] S.A. Castillo-Díaz, M.E. Garay-Sevilla, M.A. Hernández-González, M.O. Solís-Martínez, S. Zaina, Extensive demethylation of normally hypermethylated CpG islands occurs in human atherosclerotic arteries, Int. J. Mol. Med. 26 (2010) 691– 700. 10.3892/ijmm-00000515.

[13] S. Zaina, H. Heyn, F.J. Carmona, N. Varol, S. Sayols, E. Condom, J. Ramirez-Ruz, A. Gomez, I. Goncalves, S. Moran, M. Esteller, DNA Methylation Map of Human Atherosclerosis, Circ. Cardiovasc. Genet. 7 (2014) 692–700. 10.1161/CIRCGENETICS.113.000441.

[14] M. del P. Valencia-Morales, S. Zaina, H. Heyn, F.J. Carmona, N. Varol, S. Sayols, E. Condom, J. Ramírez-Ruz, A. Gomez, S. Moran, G. Lund, D. Rodríguez-Ríos, G. López-González, M. Ramírez-Nava, C. de la Rocha, A. Sanchez-Flores, M. Esteller, The DNA methylation drift of the atherosclerotic aorta increases with lesion progression., BMC Med. Genomics. 8 (2015) 7. 10.1186/s12920-015-0085-1.

[15] D. Jiang, M. Sun, L. You, K. Lu, L. Gao, C. Hu, S. Wu, G. Chang, H. Tao, D. Zhang, DNA methylation and hydroxymethylation are associated with the degree of coronary atherosclerosis in elderly patients with coronary heart disease, Life Sci. (2019). 10.1016/j.lfs.2019.03.021.

[16] C. de la Rocha, S. Zaina, G. Lund, Is Any Cardiovascular Disease-Specific DNA Methylation Biomarker Within Reach?, Curr. Atheroscler. Rep. 22 (2020) 62. 10.1007/s11883-020-00875-3.

[17] R. Rangel-Salazar, M. Wickström-Lindholm, C.A. Aguilar-Salinas, Y. Alvarado-Caudillo, K.B. V Døssing, M. Esteller, E. Labourier, G. Lund, F.C. Nielsen, D. Rodríguez-Ríos, M.O. Solís-Martínez, K. Wrobel, K. Wrobel, S. Zaina, Human native lipoprotein-induced de novo DNA methylation is associated with repression of inflammatory genes in THP-1 macrophages., BMC Genomics. 12 (2011) 582. 10.1186/1471-2164-12-582.

[18] G.A. Silva-Martínez, D. Rodríguez-Ríos, Y. Alvarado-Caudillo, A. Vaquero, M. Esteller, F.J. Carmona, S. Moran, F.C. Nielsen, M. Wickström-Lindholm, K. Wrobel, K. Wrobel, G. Barbosa-Sabanero, S. Zaina, G. Lund, Arachidonic and oleic acid exert distinct effects on the DNA methylome, Epigenetics. 11 (2016) 321–334. 10.1080/15592294.2016.1161873.

[19] E. Hall, P. Volkov, T. Dayeh, K. Bacos, T. Rönn, M.D. Nitert, C. Ling, Effects of palmitate on genome-wide mRNA expression and DNA methylation patterns in human pancreatic islets., BMC Med. 12 (2014) 103. 10.1186/1741-7015-12-103.

[20] A. Koseler, F. Ma, I.D. Kilic, M. Morselli, O. Kilic, M. Pellegrini, Genome-wide DNA Methylation Profiling of Blood from Monozygotic Twins Discordant for Myocardial Infarction., In Vivo. 34 (2020) 361–367. 10.21873/invivo.11782.

[21] Y.Z. Jiang, J.M. Jiménez, K. Ou, M.E. McCormick, L. Di Zhang, P.F. Davies, Hemodynamic disturbed flow induces differential DNA methylation of endothelial Kruppel-like factor 4 promoter in vitro and in vivo, Circ. Res. 115 (2014) 32–43. 10.1161/CIRCRESAHA.115.303883.

[22] B. Li, G. Zang, W. Zhong, R. Chen, Y. Zhang, P. Yang, J. Yan, Activation of CD137 signaling promotes neointimal formation by attenuating TET2 and transferrring from endothelial cell-derived exosomes to vascular smooth muscle cells., Biomed. Pharmacother. 121 (2019) 109593. 10.1016/j.biopha.2019.109593.

[23] Z. Zhaolin, C. Jiaojiao, W. Peng, L. Yami, Z. Tingting, T. Jun, W. Shiyuan, X. Jinyan, W. Dangheng, J. Zhisheng, W. Zuo, OxLDL induces vascular endothelial cell pyroptosis through miR-125a-5p/TET2 pathway, J. Cell. Physiol. 234 (2018) 7475–7491. 10.1002/jcp.27509.

[24] J. Yu, Y. Qiu, J. Yang, S. Bian, G. Chen, M. Deng, H. Kang, L. Huang, DNMT1-PPARγ pathway in macrophages regulates chronic inflammation and atherosclerosis development in mice., Sci. Rep. 6 (2016) 30053. 10.1038/srep30053.

[25] J. Zhuang, P. Luan, H. Li, K. Wang, P. Zhang, Y. Xu, W. Peng, The yin-yang dynamics of DNA methylation is the key regulator for smooth muscle cell phenotype switch and vascular remodeling, Arterioscler. Thromb. Vasc. Biol. 37 (2017) 84–97. 10.1161/ATVBAHA.116.307923.

[26] J. Dunn, H. Qiu, S. Kim, D. Jjingo, R. Hoffman, C.W. Kim, I. Jang, D.J. Son, D. Kim, C. Pan, Y. Fan, I.K. Jordan, H. Jo, Flow-dependent epigenetic DNA methylation regulates endothelial gene expression and atherosclerosis, J. Clin. Invest. 124 (2014) 3187–3199. 10.1172/JCI74792.(24).

[27] Q. Cao, X. Wang, L. Jia, A.K. Mondal, A. Diallo, G.A. Hawkins, S.K. Das, J.S. Parks, L. Yu, H. Shi, H. Shi, B. Xue, Inhibiting DNA methylation by 5-Aza-2’-deoxycytidine ameliorates atherosclerosis through suppressing macrophage inflammation, Endocrinology. 155 (2014). 10.1210/en.2014-1595.

[28] K.A. Strand, S. Lu, M.F. Mutryn, L. Li, Q. Zhou, B.T. Enyart, A.J. Jolly, A.M. Dubner, K.S. Moulton, R.A. Nemenoff, K.A. Koch, D. LaBarbera, M.C.M. Weiser-Evans, High Throughput Screen Identifies the DNMT1 (DNA Methyltransferase-1) Inhibitor, 5-Azacytidine, as a Potent Inducer of PTEN (Phosphatase and Tensin Homolog), Arterioscler. Thromb. Vasc. Biol. 40 (2020) 1854–1869. 10.1161/ATVBAHA.120.314458.

[29] J. Peng, Q. Yang, A.-F. Li, R.-Q. Li, Z. Wang, L.-S. Liu, Z. Ren, X.-L. Zheng, X.-Q. Tang, G.-H. Li, Z.-H. Tang, Z.-S. Jiang, D.-H. Wei, Tet methylcytosine dioxygenase 2 inhibits atherosclerosis via upregulation of autophagy in ApoE-/- mice., Oncotarget. 7 (2016) 76423–76436. 10.18632/oncotarget.13121.

[30] I. Hassanin, A. Elzoghby, Albumin-based nanoparticles: a promising strategy to overcome cancer drug resistance, Cancer Drug Resist. 3 (2020) 930–976. 10.20517/cdr.2020.68.

[31] M. Ahmed, R. Baumgartner, S. Aldi, P. Dusart, U. Hedin, B. Gustafsson, K. Caidahl, Human serum albumin-based probes for molecular targeting of macrophage scavenger receptors., Int. J. Nanomedicine. 14 (2019) 3723–3741. 10.2147/IJN.S197990.

[32] L.H. Li, E.J. Olin, H.H. Buskirk, L.M. Reineke, Cytotoxicity and Mode of Action of 5-Azacytidine on L1210 Leukemia, Cancer Res. 30 (1970).

[33] J. Veselý, A. Čihák, Incorporation of a potent antileukemic agent, 5-aza-2’-deoxycytidine, into DNA of cells from leukemic mice, Cancer Res. 37 (1977) 3684–3689. https://pubmed.ncbi.nlm.nih.gov/71199/ (accessed January 13, 2022).

[34] F.M. Segers, H. Yu, T.J. Molenaar, P. Prince, T. Tanaka, T.J. van Berkel, E.A. Biessen, Design and validation of a specific scavenger receptor class AI binding peptide for targeting the inflammatory atherosclerotic plaque., Arterioscler. Thromb. Vasc. Biol. 32 (2012) 971–978. 10.1161/ATVBAHA.111.235358.

[35] A. Sánchez-Arreguin, R. Carriles, N. Ochoa-Alejo, M.G. López, L. Sánchez-Segura, Generation of BSA-capsaicin Nanoparticles and Their Hormesis Effect on the Rhodotorula mucilaginosa Yeast, Molecules. 24 (2019). 10.3390/molecules24152800.

[36] C. Chilom, G. BaranL, D. Gǎzdaru, A. Popescu, Characterisation by fluorescence of human and bovine serum albumins in interaction with eosin Y, J. Optoelectron. Adv. Mater. 15 (2013).

[37] S.H. Zhang, R.L. Reddick, J.A. Piedrahita, N. Maeda, Spontaneous hypercholesterolemia and arterial lesions in mice lacking apolipoprotein E., Science. 258 (1992) 468–471. http://www.ncbi.nlm.nih.gov/pubmed/1411543 (accessed April 7, 2015).

[38] L. Brånén, L. Pettersson, M. Lindholm, S. Zaina, A procedure for obtaining whole mount mouse aortas that allows atherosclerotic lesions to be quantified., Histochem. J. 33 (2001) 227–229. http://www.ncbi.nlm.nih.gov/pubmed/11550804 (accessed May 6, 2015).

[39] E. Cohen-Sela, D. Dangoor, H. Epstein, I. Gati, H.D. Danenberg, G. Golomb, J. Gao, Nanospheres of a bisphosphonate attenuate intimal hyperplasia., J. Nanosci. Nanotechnol. 6 (2006) 3226–3234. http://www.ncbi.nlm.nih.gov/pubmed/17048541 (accessed July 20, 2017).

[40] J. Yoo, S. Choi, J.L. Medina-Franco, Molecular modeling studies of the novel inhibitors of DNA methyltransferases SGI-1027 and CBC12: implications for the mechanism of inhibition of DNMTs., PLoS One. 8 (2013) e62152. 10.1371/journal.pone.0062152.

[41] J. Datta, K. Ghoshal, W.A. Denny, S.A. Gamage, D.G. Brooke, P. Phiasivongsa, S. Redkar, S.T. Jacob, A New Class of Quinoline-Based DNA Hypomethylating Agents Reactivates Tumor Suppressor Genes by Blocking DNA Methyltransferase 1 Activity and Inducing Its Degradation, Cancer Res. 69 (2009) 4277–85. 10.1158/0008-5472.CAN-08-3669.

[42] N. Sun, J. Zhang, C. Zhang, B. Zhao, A.O. Jiao, DNMTs inhibitor SGI-1027 induces apoptosis in Huh7 human hepatocellular carcinoma cells, Oncol. Lett. 16 (2018). 10.3892/ol.2018.9390.

[43] M.J. Mitchell, M.M. Billingsley, R.M. Haley, M.E. Wechsler, N.A. Peppas, R. Langer, Engineering precision nanoparticles for drug delivery, Nat. Rev. Drug Discov. 20 (2020). 10.1038/s41573-020-0090-8.

[44] M. Cooley, A. Sarode, M. Hoore, D.A. Fedosov, S. Mitragotri, A. Sen Gupta, Influence of particle size and shape on their margination and wall-adhesion: implications in drug delivery vehicle design across nano-to-micro scale., Nanoscale. 10 (2018) 15350–15364. 10.1039/c8nr04042g.

[45] Q. Cao, X. Wang, L. Jia, A.K. Mondal, A. Diallo, G.A. Hawkins, S.K. Das, J.S. Parks, L. Yu, H. Shi, H. Shi, B. Xue, Inhibiting DNA Methylation by 5-Aza-2’-deoxycytidine ameliorates atherosclerosis through suppressing macrophage inflammation., Endocrinology. 155 (2014) 4925–4938. 10.1210/en.2014-1595.

[46] M.A. Zimmerman, L.L. Reznikov, C.D. Raeburn, C.H. Selzman, Interleukin-10 attenuates the response to vascular injury, J. Surg. Res. 121 (2004). 10.1016/j.jss.2004.03.025.

[47] E. Galkina, K. Ley, Immune and inflammatory mechanisms of atherosclerosis (*), Annu Rev Immunol. 27 (2009) 165–197. 10.1146/annurev.immunol.021908.132620.

[48] Z. Mallat, S. Besnard, M. Duriez, V. Deleuze, F. Emmanuel, M.F. Bureau, F. Soubrier, B. Esposito, H. Duez, C. Fievet, B. Staels, N. Duverger, D. Scherman, A. Tedgui, Protective role of interleukin-10 in atherosclerosis, Circ Res. 85 (1999) e17–24.

[49] J.L. Stoger, P. Goossens, M.P. de Winther, Macrophage heterogeneity: relevance and functional implications in atherosclerosis, Curr Vasc Pharmacol. 8 (2010) 233–248. https://doi.org/BSP/CVP/E-Pub/000067 [pii].

[50] G.K. Hansson, P. Libby, The immune response in atherosclerosis: a double-edged sword, Nat Rev Immunol. 6 (2006) 508–519. https://doi.org/nri1882 [pii]10.1038/nri1882.

[51] A. Hermansson, D.F. Ketelhuth, D. Strodthoff, M. Wurm, E.M. Hansson, A. Nicoletti, G. Paulsson-Berne, G.K. Hansson, Inhibition of T cell response to native low-density lipoprotein reduces atherosclerosis, J Exp Med. 207 (2010) 1081– 1093. https://doi.org/jem.20092243 [pii]10.1084/jem.20092243.

[52] S. Poirier, S. Samami, M. Mamarbachi, A. Demers, T.Y. Chang, D.E. Vance, G.M. Hatch, G. Mayer, The epigenetic drug 5-azacytidine interferes with cholesterol and lipid metabolism., J. Biol. Chem. 289 (2014) 18736–18751. 10.1074/jbc.M114.563650.

[53] F.L.T. Gomes, R.C. Maranhão, E.R. Tavares, P.O. Carvalho, M.L. Higuchi, F.R. Mattos, F.G. Pitta, S.A. Hatab, R. Kalil-Filho, C. V. Serrano, Regression of Atherosclerotic Plaques of Cholesterol-Fed Rabbits by Combined Chemotherapy With Paclitaxel and Methotrexate Carried in Lipid Core Nanoparticles, J. Cardiovasc. Pharmacol. Ther. 23 (2018). 10.1177/1074248418778836.

[54] A.A. Manolis, T.A. Manolis, H. Melita, D.P. Mikhailidis, A.S. Manolis, Low serum albumin: A neglected predictor in patients with cardiovascular disease., Eur. J. Intern. Med. 102 (2022) 24–39. 10.1016/j.ejim.2022.05.004.

[55] X. Zhong, H. Jiao, D. Zhao, J. Teng, Association between serum albumin levels and paroxysmal atrial fibrillation by gender in a Chinese population: a case-control study., BMC Cardiovasc. Disord. 22 (2022) 387. 10.1186/s12872-022-02813-4.

[56] R.F. Gillum, The association between serum albumin and HDL and total cholesterol., J. Natl. Med. Assoc. 85 (1993) 290–292.

