## Supplemental Figures for "Macrophage-targeted DNA methyltransferase inhibitor SGI-1027 decreases atherosclerosis in ApoE-null mice"

A

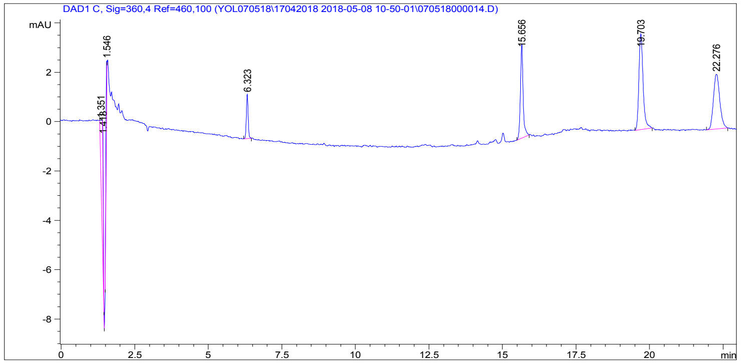


Signal 1: DAD1 C, Sig=360,4 Ref=460,100

| RetTime [min] | Type | Area [mAU*s] | Amt/Area | Amount [mg/ml] | Grp | Name |
|---|---|---|---|---|---|---|
| 6.323 | BB | 7.55517 | 0.00000 | 0.00000 | | SGI |

Totals : 0.00000

B

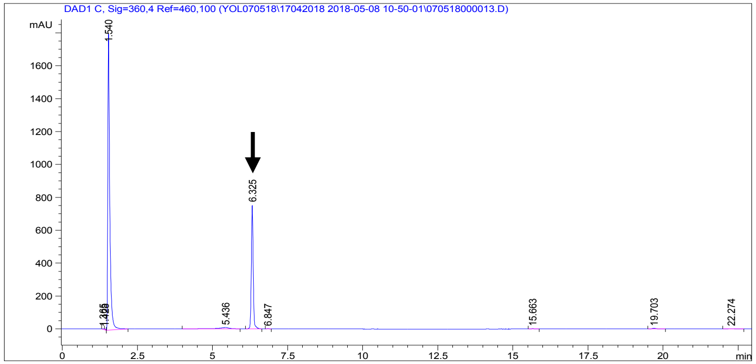


Signal 1: DAD1 C, Sig=360,4 Ref=460,100

| RetTime [min] | Type | Area [mAU*s] | Amt/Area | Amount [mg/ml] | Grp | Name |
|---|---|---|---|---|---|---|
| 6.325 | BB | 3269.34546 | 2.98388e-5 | 9.75532e-2 | | SGI |

Totals : 9.75532e-2

**Supplementary Figure 1. HPLC detection of NP-loaded SGI-1027.** Representative HPLC chromatograms of NP ethanolic extracts. A, void NP. B, SGI-1027-loaded, FLAGPP1-functionalized NP. The arrow indicates the SGI-1027 peak.

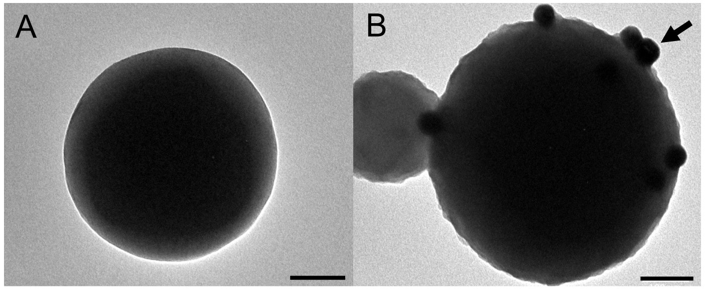


**Supplementary Figure 2. Immunogold TEM analysis of HSANP surface functionalization.** A, unfunctionalized NP. B, FLAGPP1-functionalized NP. The arrow indicates a labelled FLAGPP1 peptide. Bar, 100 nm.

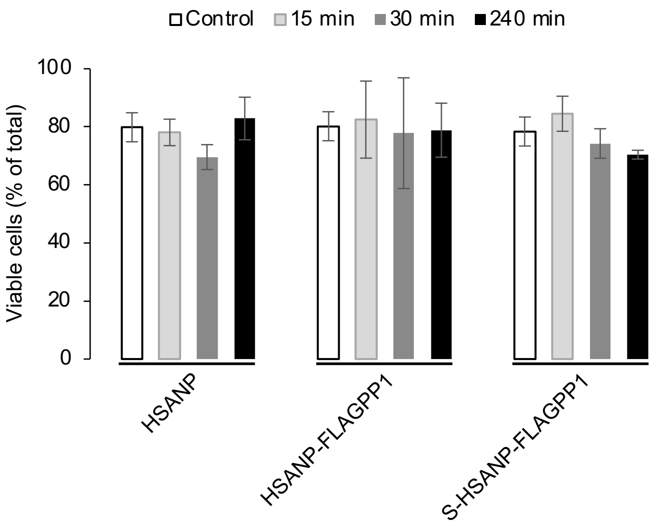


**Supplementary Figure 3. Viability of NP-challenged THP-1 macrophages.** Length of challenge with NP is indicated above the graph. n=3. Data are average±SD. HSANP, void NP; HSANP-FLAGPP1, functionalized void NP; S-HSANP-FLAGPP1, SGI-1027-loaded and functionalized NP.

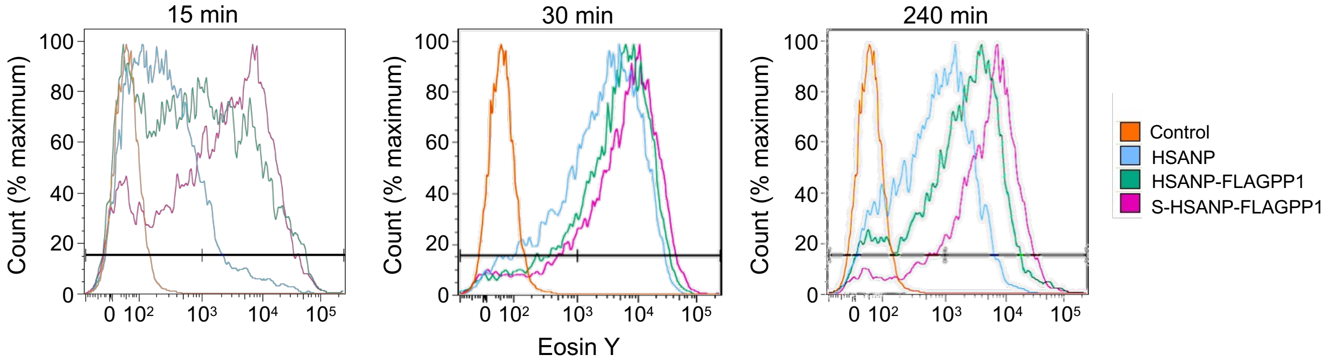


**Supplementary Figure 4. NP internalization in THP-1 macrophages.** Count/fluorescence plots of flow cytometry analysis of cell-associated eosin Y fluorescence at the indicated length of challenge with different NP, each represented by the colour code indicated on the right. Time zero control is used as reference in all plots. HSANP, void NP; HSANP-FLAGPP1, functionalized NP; S-HSANP-FLAGPP1, SGI-1027-loaded and functionalized NP.

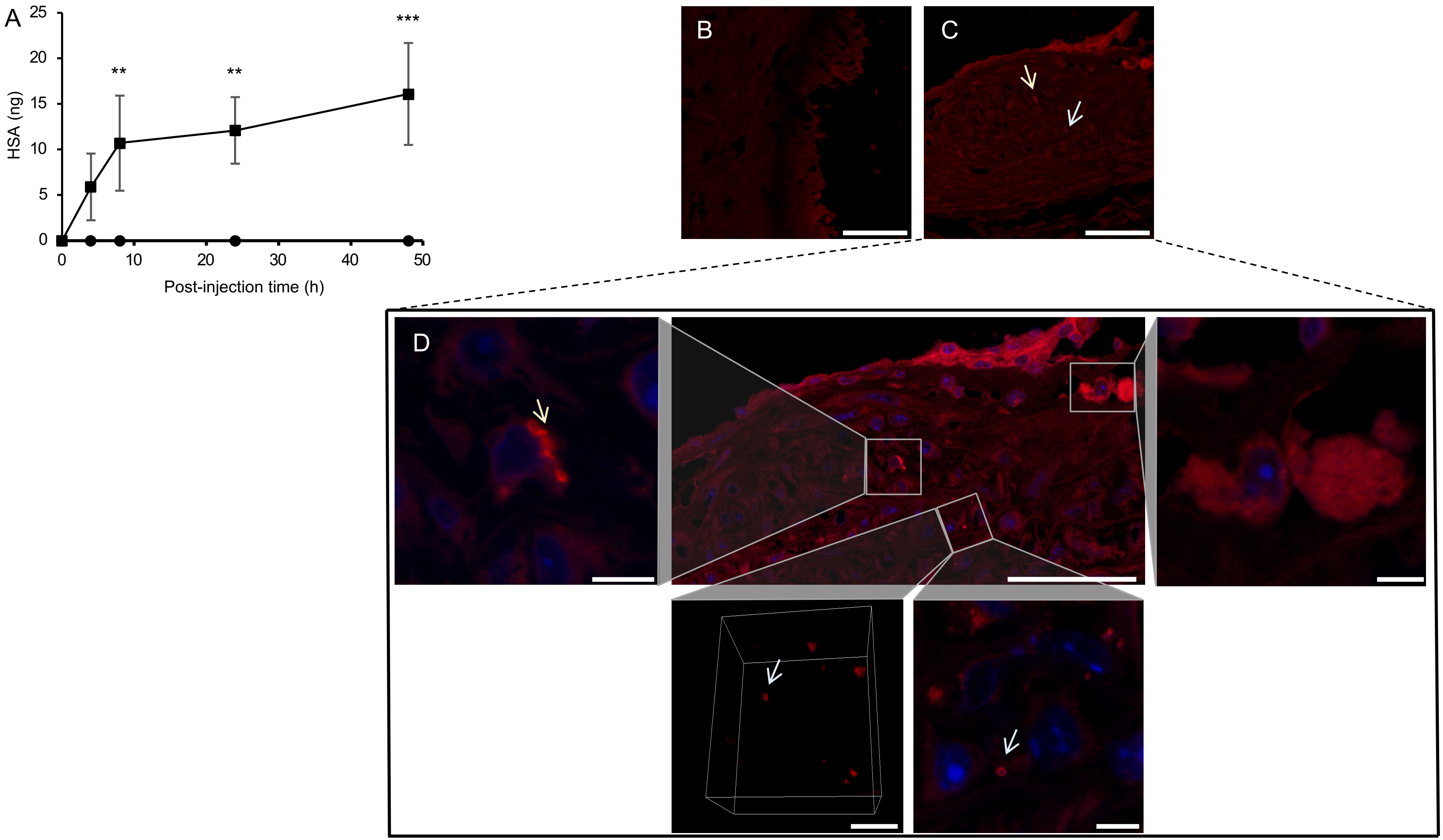


**Supplementary Figure 5. Subcutaneously injected HSANP colonize the peripheral circulation and the aortic root lesion.** A, time-course of detectable HSANP in mouse peripheral blood after subcutaneous injection. Squares and circles, HSANP-injected and PBS (vehicle)-injected mice, respectively (n=3 each). Significance is shown for comparisons with the 4 h time point. ANOVA and Bonferroni *post hoc* test. B-D, confocal images of immunogold-labelled FLAGPP1. B, vehicle-treated mice. C, S-HSANP-FLAGPP1-treated mice. D, zoomed-in details of the Z-stack for the regions of interest, including a 3D projection of one intracellular signal. Arrows indicate the dark red signal above autofluorescence (B). Notice the immunogold signal at the edge of and within the atheroma. Bar in B,C is 50 μm. Bar in zoom-in pictures is 5 μm.

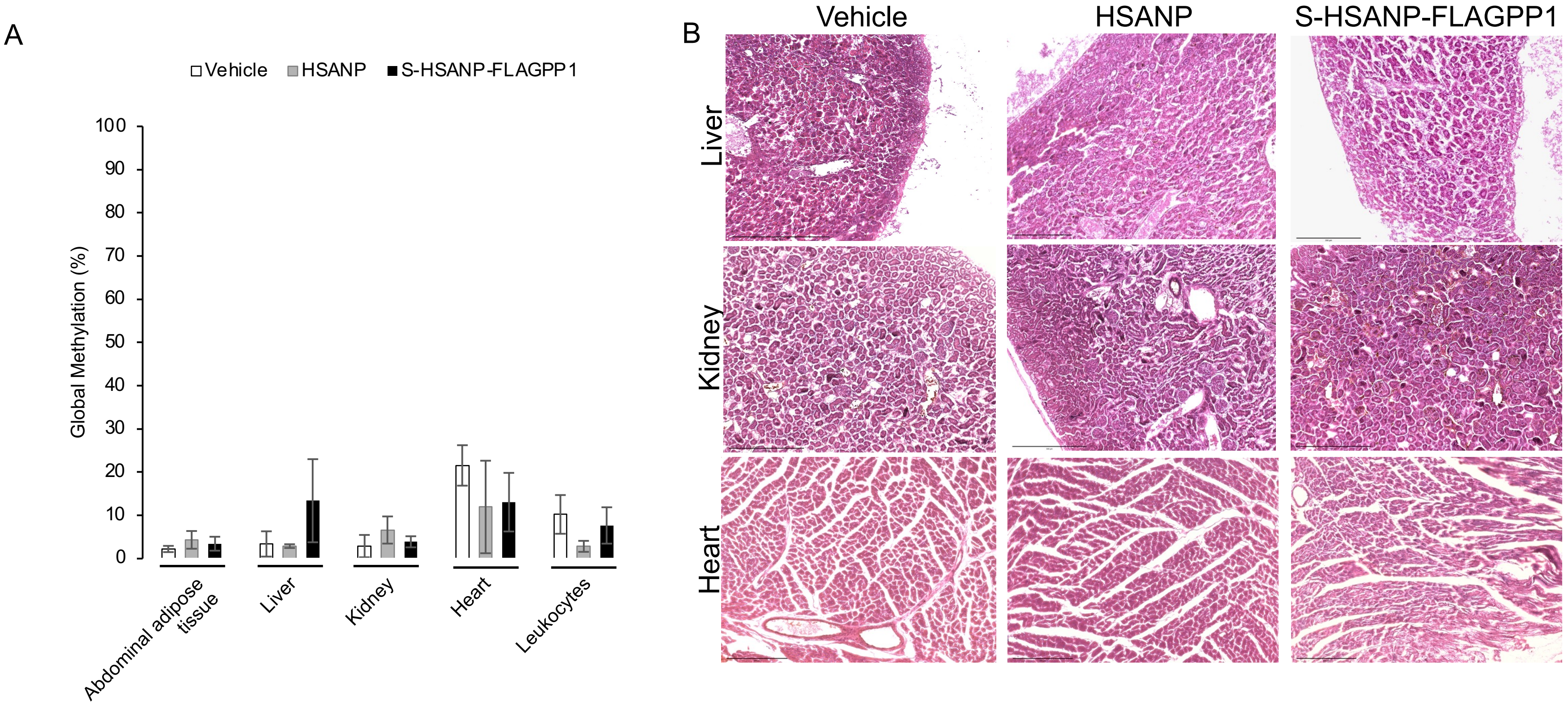


**Supplementary Figure 6. Effects of NP on non-target tissues.** A, global DNA methylation levels of selected non-vascular tissues (n=3/group). B, representative HE staining of liver, kidney and heart. Bar, 200 nm. HSANP, void NP; S-HSANP-FLAGPP1, SGI-1027-loaded and functionalized NP. Kruskal-Wallis test.
